# Insights into substrate binding and utilization by hyaluronan synthase

**DOI:** 10.1101/2025.10.17.683186

**Authors:** Zachery Stephens, Julia Karasinska, Jochen Zimmer

## Abstract

Hyaluronan (HA), a heteropolysaccharide of alternating N-acetylglucosamine (GlcNAc) and glucuronic acid (GlcA), is an essential component of the vertebrate extracellular matrix. HA biosynthesis proceeds via three evolutionarily convergent reaction mechanisms, catalyzed by the membrane integrated class 1 enzymes that either elongate the non-reducing (NR) or reducing end of HA, as well as the class 2 hyaluronan synthase (HAS), a soluble non-processive enzyme. Class 1-NR HAS, found in both vertebrates and large double-stranded DNA viruses, is monomeric and couples HA polymerization via coordinated transfer of UDP-GlcNAc and UDP-GlcA substrates with the secretion of the nascent HA chain through its own transmembrane channel. How this HAS discriminates between two UDP-sugars using a single active site is a critical, yet unresolved question. Using single-particle cryo-EM, we reveal a two-step process by which the *Chlorella* virus HAS (CvHAS) recognizes and positions its substrate, UDP-GlcA, for glycosyl transfer. Further, we report greatly diminished turnover of UDP-GlcA in the absence of a primer, distinguishing acceptor free activity toward UDP-GlcNAc. Lastly, prompted by observation of a dodecyl maltoside bound HAS, we demonstrate the utility of non-canonical acceptors in priming of a UDP-GlcA transfer reaction. Altogether, this work clarifies the molecular basis for HAS’ dual substrate specificity and the role of UDP-GlcA recognition in integrity of HA synthesis.

## Introduction

Hyaluronan (HA), a heteropolysaccharide of alternating glucuronic acid (GlcA) and N-acetylglucosamine (GlcNAc), is an abundant and essential extracellular matrix material in vertebrates. It performs a plethora of developmental and physiological functions with critical contributions to fertilization, cardio- and angiogenesis, wound healing, and joint lubrication^1–3^.

Biosynthesis of HA is catalyzed in a cation dependent polymerization reaction with nucleotide-sugar substrates, UDP-GlcNAc and UDP-GlcA, by HA synthase (HAS)^4^. Several unique mechanisms of HA synthesis have emerged, lending to the definition of processive class 1 HASs, transmembrane proteins which couple HA synthesis and transport activities, and distributive class 2 HASs, peripheral membrane proteins which lack a secretion function^5^. HA polymerization by class 1 HAS can occur at either the non-reducing end (class 1-NR) as reported for the *Paramecium bursaria Chlorella virus* HAS, or the reducing end (class 1-R) in the case of the dimeric *S. equismillis* HAS^6^. Class 1-NR HASs function as monomers and contain a single catalytic GT-A domain^7^, necessarily dictating that a single active site be sufficient to sequentially recognize and transfer both UDP-GlcA and UDP-GlcNAc to a nascent HA chain^8^.

Recent structural and biochemical analyses of Chlorella virus and *Xenopus laevis* isoform-1 HAS (CvHAS and XlHAS-1, respectively) provided detailed insights into the multitasking of type 1-NR HAS^8–11^. First, to initiate HA biosynthesis, HAS binds and hydrolyzes UDP-GlcNAc, such that the released GlcNAc monosaccharide can prime polymer biosynthesis. Second, the GlcNAc-primed enzyme binds the next substrate, UDP-GlcA, leading to the formation of a ternary complex that facilitates glycosyl transfer and the formation of a β-linked GlcAβ1-3GlcNAc disaccharide. The subsequent binding of a UDP-GlcNAc substrate molecule presumably translocates the HA disaccharide by an unknown mechanism and facilitates the transfer of GlcNAc to the nascent HA polymer, thereby forming a β-1,4 linkage to GlcA. These steps must be repeated thousands of times to synthesize and secrete an HA polysaccharide of greater than 10,000 disaccharide units^12^.

While UDP-GlcNAc is a common metabolite in all reported HA-producing species, the physiological concentration of UDP-GlcA, however, may limit HA biosynthesis under certain conditions^13^. Accordingly, the Chlorella virus encodes a UDP-glucose dehydrogenase enzyme that generates UDP-GlcA from UDP-glucose^14^. Further, expressing this enzyme in engineered HA-producing systems increased the overall production levels^15,16^.

In this study, we delineate differences in UDP-GlcA coordination by CvHAS using cryo-electron microscopy (cryo-EM). Enzymology and biochemical analysis were utilized to understand how UDP-GlcA interaction strength and turnover efficiency vary in the presence and the absence of a GlcNAc primer. We further show that CvHAS exhibits a degree of promiscuity toward acceptors for glycosyl transfer, enabling the biosynthesis of unnatural complex carbohydrates. Lastly, our structural analysis reveals a dodecyl maltoside-inhibited state of CvHAS in which the detergent molecule occupies the acceptor-binding site, thereby providing insights into HAS substrate promiscuity.

## Results

We used cryo-EM analysis to gain structural insights into the interaction of CvHAS with its substrate UDP-GlcA. The catalytically inactive (D302N) CvHAS mutant was reconstituted into MSPE3D1 lipid nanodiscs as described before^8^ and complexed with high affinity nanobodies that recognize cytosolic and extracellular epitopes. This complex was then incubated with 10 mM MnCl_2_ and 5 mM UDP-GlcA in the absence of a receiving GlcNAc monosaccharide, prior to cryo grid preparation. The obtained cryo EM dataset was processed as outlined in Fig. S1.

### CvHAS binds UDP-GlcA in two different binding poses

Focused refinement of particles harboring a UDP-GlcA molecule at the active site resolved two substrate binding poses, interpreted as proofreading and inserted states. The inserted pose resembles the previously reported UDP-GlcA conformation in the presence of a priming GlcNAc sugar^11^. In this state, the nucleotide’s uracyl moiety is surrounded by Tyr91, His174, and Lys177, its diphosphate group, together with CvHAS’ DxD motif (Asp201 and Asp203), coordinates a divalent cation, most likely Mn^2+^, and the glucuronic acid donor sugar resides in a pocket underneath the acceptor binding site (Fig 1B). GlcA’s ring oxygen resides within hydrogen bonding distance to the Nε of Trp342, and its C6 carboxylate group, although poorly resolved, is positioned near the guanidinium group of Arg341 (Fig 1C and D). The side chain’s Nε is within 3.5 Å of GlcA’s carboxyl group, potentially accounting for a weak electrostatic interaction. An alternative side chain conformation for Arg341 was well-supported by cryo-EM density and thus modeled. In this position, the residue’s guanidinium group is about 4.6 Å from GlcA’s C6 carboxylate (Fig 1D) and likely interacts with it via a mediating water molecule. In addition, the conserved Asp201 forms a salt bridge with Lys177 at the back of the nucleotide binding pocket (Fig. 1C).

**Figure 1:**
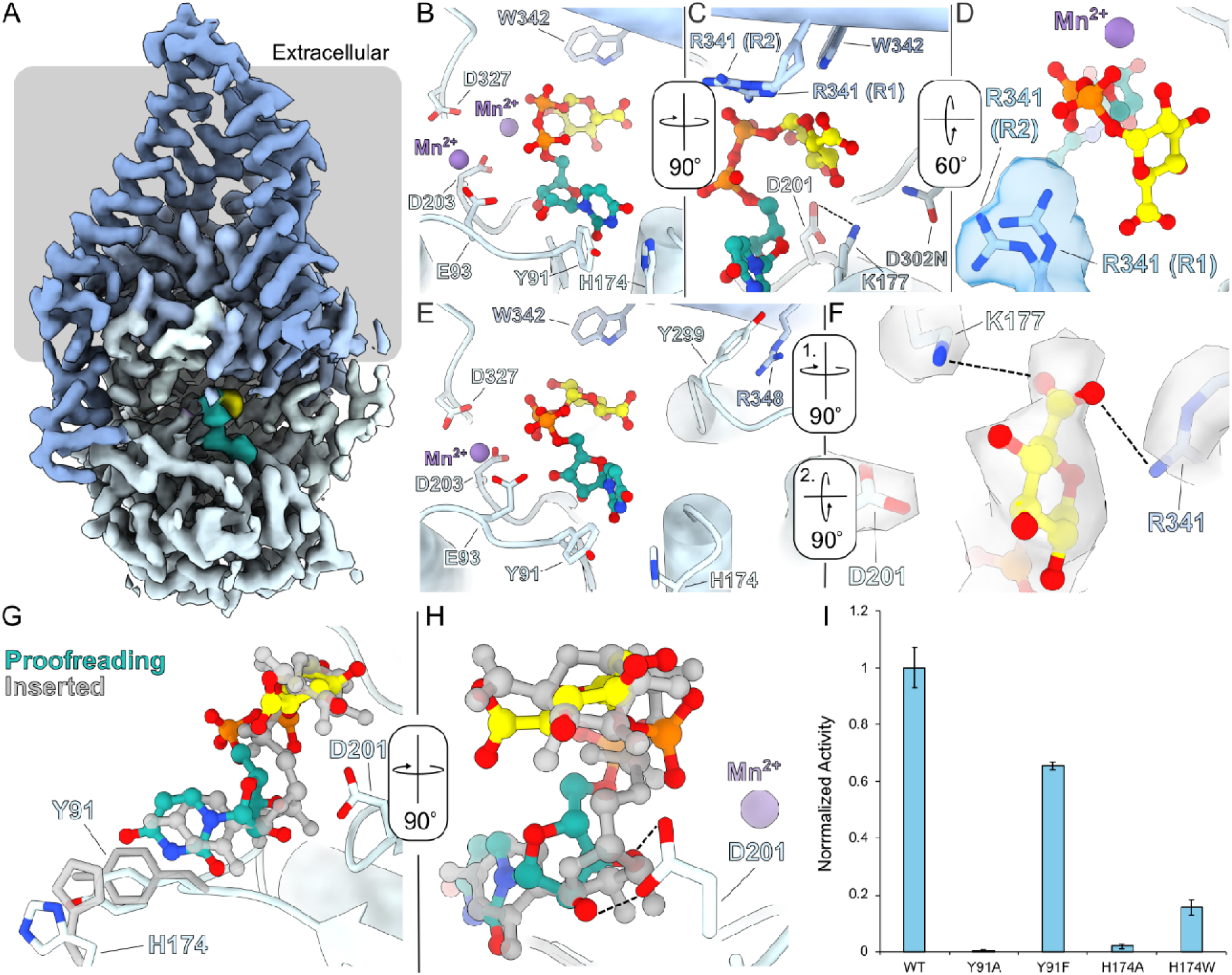
UDP-GlcA coordination and proofreading. A, CryoEM density map of CvHAS bound to UDP-GlcA with a grey bar representing the membrane boundary. UDP-GlcA density shown in light sea green and yellow for its UDP and GlcA moieties, respectively. B-C, Coordination of an inserted UDP-GlcA substrate in the absence of a GlcNAc primer. UDP-GlcA is shown as a ball and stick model with light sea green carbon atoms for the UDP moiety and yellow carbon atoms for GlcA. D, CryoEM density for Arg341 rotamer 1 (R1) and rotamer 2 (R2). E, Coordination of UDP-GlcA in the proofreading conformation. F, CryoEM density for UDP-GlcA’s sugar ring and surrounding residues in the proofreading state. G-H, Alignment of proofreading and inserted UDP-GlcA positions. The ligand in the inserted UDP-GlcA structure is shown in grey. H, Activity of uracil binding pocket mutants for CvHAS. Activity measurements were normalized to wild-type (WT), and were reported as the average of three technical replicates. Error bars represent the standard deviation from the mean.

Previous analysis identified a second divalent cation binding site in CvHAS’ catalytic pocket^8^. This site is created by Glu93 together with the C-terminal Asp of the DxD motif (Asp203) and appears to be metal occupied in nucleotide-bound and apo states. This is also the case in both of our new cryo EM maps, suggesting that the site is sufficient to coordinate a hydrated cation (Fig S2A-D). Further, this coordination site is required for catalytic activity as replacing Glu93 with Ala renders CvHAS inactive ^8^.

In the second ‘proofreading’ UDP-GlcA binding pose resolved in our dataset, the ligand’s nucleotide and donor sugar moieties are less deeply inserted into the catalytic pocket (Fig. 1E-H). The uracil moiety is shifted by about 1.5 Å towards the entrance of the catalytic pocket, resulting in the rotation of His174 away from Tyr91. Further, the nucleotide’s ribose moiety is tilted away from the DxD motif, which places the attached diphosphate group also closer to the binding cleft’s entrance. In this position, the diphosphate is suitably positioned to contribute to cation coordination at the second metal binding site, together with Asp203 and Glu93 (Fig 1E). Indeed, we observe strong cryo EM density at this site, consistent with an octahedrally coordinated manganese cation (Fig S2C and S2D).

The partial insertion of UDP-GlcA in the proofreading pose further allows Asp201 of the DxD motif to move away from its binding partner Lys177 to interact with the ribose’s C2 and C3 hydroxyl groups (Fig 1H). The ammonium group of Lys177, instead, moves towards the donor sugar and, together with the guanidinium group of Arg341, sandwiches GlcA’s carboxylate group (Fig 1F). Both side chains are well resolved in the cryo EM map and about 3.5 Å away from the carboxylate group, which is also resolved at a slightly lower contour level (Fig 1F). Compared to the inserted UDP-GlcA state described above, these interactions create a basic pocket that recognizes GlcA’s carboxylate group.

Next to the interactions with Lys177 and Arg341, GlcA’s carboxylate is further framed by the C-terminal segment of the priming loop that leads into the conserved GDD motif (residues 300-302). The backbone conformation of the priming loop differs slightly in both UDP-GlcA bound conformations. Most notably, while the entire backbone is well resolved in the inserted UDP-GlcA bound pose (Fig S2E), the density of the conserved Gly300 is essentially absent in the proofreading state (Fig S2F), suggesting that this residue and parts of the preceding priming loop are flexible until UDP-GlcA is fully inserted into the catalytic pocket. The conserved Arg348 that belongs to an amphipathic helix at the cytosolic water-lipid interface, interacts with the backbone carbonyl oxygen of Gly300 in the inserted state. When the priming loop is flexible in the proofreading state, however, Arg348 bends away by about 2 Å to adopt a cation-π stacking interaction with Tyr299 (Fig. 1E). Arginine 348 is conserved among bacterial and eukaryotic HASs, and Tyr299 is conservatively substituted with Phe in vertebrate HASs, suggesting this interaction is preserved across species.

We next thought to validate the contribution of conformational changes in the uracil binding groove to catalytic activity of CvHAS. To this end, we tested the functional relevance of Tyr91 and His174 through site-directed mutagenesis. CvHAS’ activity can be quantified in vitro by measuring the accumulation of tritiated HA by scintillation counting, as previously described^6^. Accordingly, replacing Tyr91 with Ala abolishes catalytic activity of CvHAS, while substituting the residue with Phe maintains about 80% activity, relative to the wild-type enzyme (Fig 1I). Similarly drastic effects are observed when replacing His174 with either Ala or Trp, its corresponding substitution in vertebrate HASs. The Ala mutant is essentially inactive while the H174W mutant retains about 25% of wild-type activity (Fig 1I). All mutants share similar size exclusion chromatography profiles with the WT enzyme, suggesting that the substitutions do not cause a folding defect (Fig. S3).

### CvHAS binds UDP-GlcA with low micromolar affinity

We performed isothermal titration calorimetry to determine the apparent affinity of UDP-GlcA binding to CvHAS. Titrating UDP-GlcA into a cell containing the catalytically inactive D302N CvHAS mutant at 45 µM revealed saturable exothermic heat responses that could be fit to a single binding isotherm. By this method, we obtained dissociation constants (K_d_) for UDP-GlcA of 69 and 24 µM in the absence and the presence of a GlcNAc primer, respectively (Fig 2A and 2B). Under similar conditions, no binding was observed for UDP-GlcNAc, UDP alone, as well as UDP-glucose (Fig S4). While the lack of a detectable heat signal for UDP-GlcNAc binding is unknown, the apparent lack of UDP and UDP-glucose binding suggests the contribution of UDP-GlcA’s carboxylate to binding.

**Figure 2:**
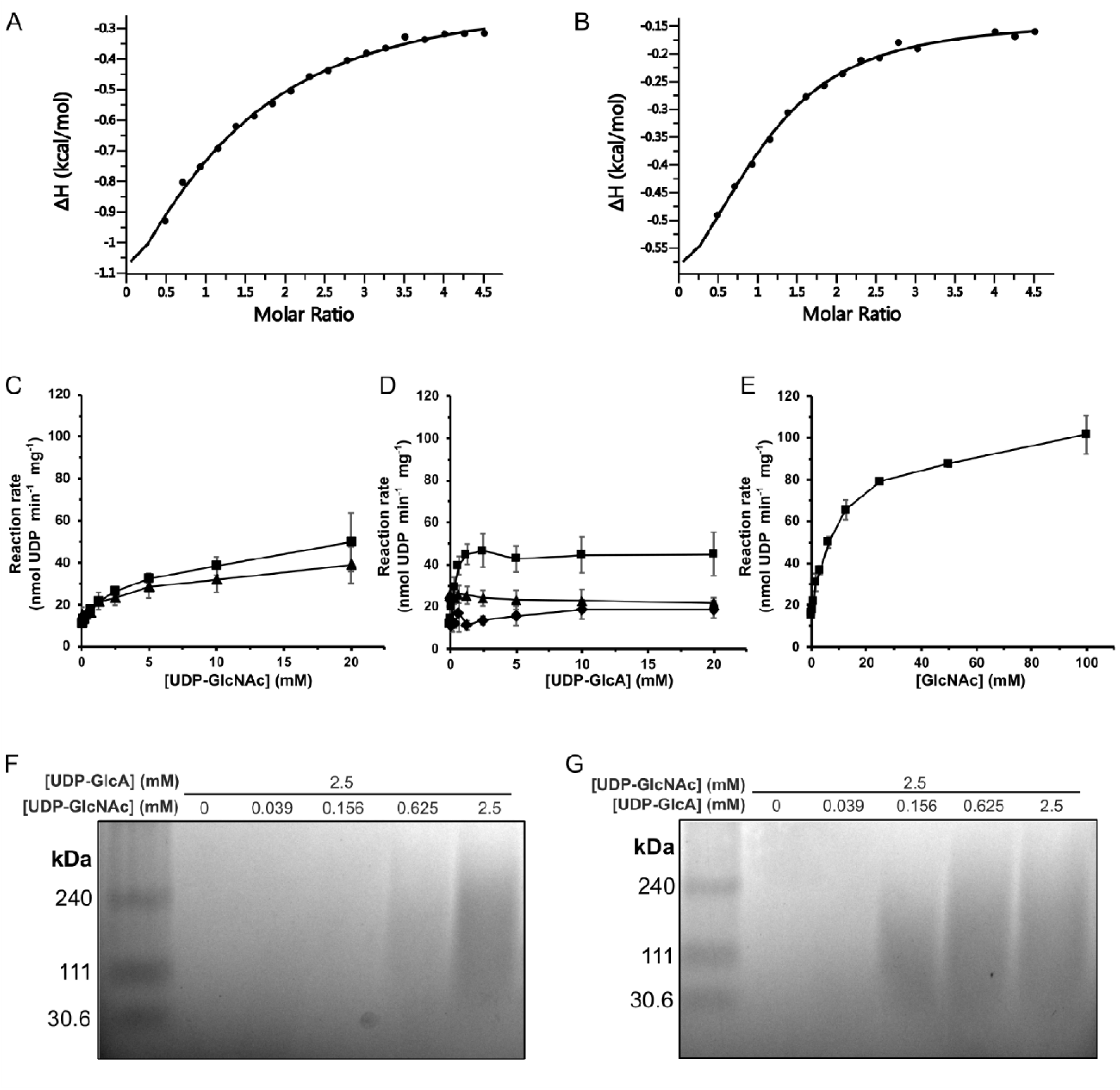
Biochemical analysis of UDP-GlcA and UDP-GlcNAc utilization by CvHAS. A-B, Binding isotherms derived from ITC experiments where CvHAS was titrated with UDP-GlcA in the absence (A) or presence (B) of GlcNAc. C, Scatter plot of UDP-GlcNAc turnover without an acceptor (▪) or in the presence of 2.5 mM UDP-GlcA (▴). D, UDP-GlcA turnover without an acceptor (◆), in the presence of 10 mM GlcNAc (▪) and in the presence of 2.5 mM UDP-GlcNAc (▴). Data points for panels C and D are reported as the average of five technical replicates. E, Measurement of UDP-GlcA turnover at a constant concentration of 2.0 mM with titration of GlcNAc (▪). Data points were reported as the average of three technical replicates. F, HA electrophoresis gel analyzing HA production under UDP-GlcNAc limiting conditions with excess UDP-GlcA. All error bars represent the standard deviation from the mean. G, HA electrophoresis gel analyzing HA production under UDP-GlcA limiting conditions with excess UDP-GlcNAc. Confirmation of HA product identity by HA lyase digestion under the provided synthesis conditions has been described in previous reports^10^. Molecular weight standards correspond to a Select-HA LoLadder (Echelon Biosciences).

We next thought to compare substrate turnover efficiency for UDP-GlcA and UDP-GlcNAc under initiation and synthesis conditions. To this end, an enzyme-coupled assay was utilized that links the release of UDP upon glycosyl transfer to the activities of pyruvate kinase and lactate dehydrogenase, as previously described^6,17^. First, measurements of UDP-GlcA and UDP-GlcNAc turnover in the absence of an acceptor substrate were performed. We observed an incremental increase in reaction rate in response to UDP-GlcNAc titration, allowing us to derive an apparent K_m_ of 744 μM. (Fig 2C and S5A).

Second, when titrating UDP-GlcA, no apparent change in substrate turnover rate above background was observed (Fig 2D), as also previously described^8^. This suggests that CvHAS is unable to hydrolyze UDP-GlcA.

Third, a kinetic experiment reflecting the first glycosyl transfer reaction performed by CvHAS was setup. Here, excess GlcNAc was supplemented to generate a primed state and UDP-GlcA was titrated. Under those conditions, a steep kinetic response to substrate titration was observed, with an apparent K_m_ for UDP-GlcA of 147 μM (Fig 2D and S5B).

Fourth, to compare kinetics of substrate turnover during initiation and processive HA synthesis, a separate set of experiments was performed. First, UDP-GlcNAc was titrated in the presence of excess UDP-GlcA. Here, maximum reaction velocity was approximately 25% lower than that measured in the presence of UDP-GlcNAc only, however the substrate concentration at which saturation was reached appeared unchanged (Fig. 2C). Second, titration of UDP-GlcA at an excess of UDP-GlcNAc revealed a constant or even slightly declining substrate turnover rate (Fig 2D). This finding suggests that the UDP-GlcNAc turnover rate remains the same, regardless of whether GlcA or presumably water forms the acceptor of the glycosyl transfer reaction.

To confirm that HA was synthesized under the titration conditions, we visualized the HA product by gel electrophoresis and ‘Stains-All’ staining^11^. To this end, synthesis reactions were quenched after 1.5 hours and electrophoresed through an agarose gel, as previously described^11^. Previous work showed that HA made by CvHAS appears as a polydisperse distribution in a molecular weight range between 30-300 kDa. Our HA electrophoresis experiment confirmed formation of a similar product, with HA signal initially observed at a UDP-GlcNAc concentration of 625 μM (Fig 2F), and a UDP-GlcA concentration of 156 μM (Fig 2G).

### GlcNAc primer affinity

Turnover of UDP-GlcA by CvHAS requires a GlcNAc acceptor, either in the form of a primer or the non-reducing end terminal HA moiety. The enzyme-coupled assay described above allowed us to determine the apparent affinity of CvHAS for the GlcNAc primer. While only minimal UDP-GlcA hydrolysis by CvHAS is observed in the absence of a GlcNAc primer^8^, CvHAS shows a drastic increase in UDP-GlcA consumption in the presence of saturating GlcNAc. This is likely due to the supplemented monosaccharide serving as the acceptor. Accordingly, titrating GlcNAc into a reaction of CvHAS at a constant UDP-GlcA concentration of 2 mM revealed a saturable increase in UDP-GlcA turnover with an apparent K_m_ of 4.0 mM (Fig 2E and S5C). We note that the apparent maximum catalytic rate observed in the presence of saturating GlcNAc is about twice the rate observed in the presence of saturating UDP-GlcA and UDP-GlcNAc (Fig. 2C and E). While the reaction in the presence of the GlcNAc primer (Fig. 2E) likely generates disaccharides that diffuse away from the enzyme, the formation of a proper HA polymer in the presence of both UDP-activated substrates (Fig. 2C) appears to reduce CvHAS’ overall catalytic rate.

### Detergent interactions at the acceptor site

Structural and functional analyses of CvHAS are routinely performed in lipid nanodiscs or the detergent glyco-diosgenin (GDN)^8^. Both conditions support the catalytic activity of the enzyme. However, the protein is initially extracted from membranes in the detergent dodecyl-β-D-maltopyranoside (DDM) that is later removed during the purification procedure (see Methods). In a DDM-solubilized state, CvHAS and XlHAS-1 are catalytically inactive (Fig S6), yet regain activity after nanodisc reconstitution or exchange into GDN detergent.

To our surprise, careful analysis of a cryo EM dataset in which CvHAS was supplemented with 2 mM MnCl_2_ and 1 mM UDP-GlcA failed to converge on a substrate-bound map. However, approximately 50% of all particles that generated a high-resolution map retained a well-resolved DDM molecule near the active site (Fig 3A and S7). The first glucosyl unit of the maltoside moiety stacks against Trp342 at the acceptor site. The second non-reducing glucosyl unit protrudes into the nucleotide binding cleft, thereby overlapping with the donor sugar binding site (Fig 3B). DDM’s interaction with Trp342 is intriguing because this residue also stabilizes the GlcNAc monosaccharide in the primed state^8^ or the terminal GlcNAc unit in an HA associated state^11^. Indeed, the maltose moiety occupies a position similar to the previously described HA disaccharide binding pose obtained after extending a GlcNAc primer with GlcA^11^ (Fig. 3C).

**Figure 3:**
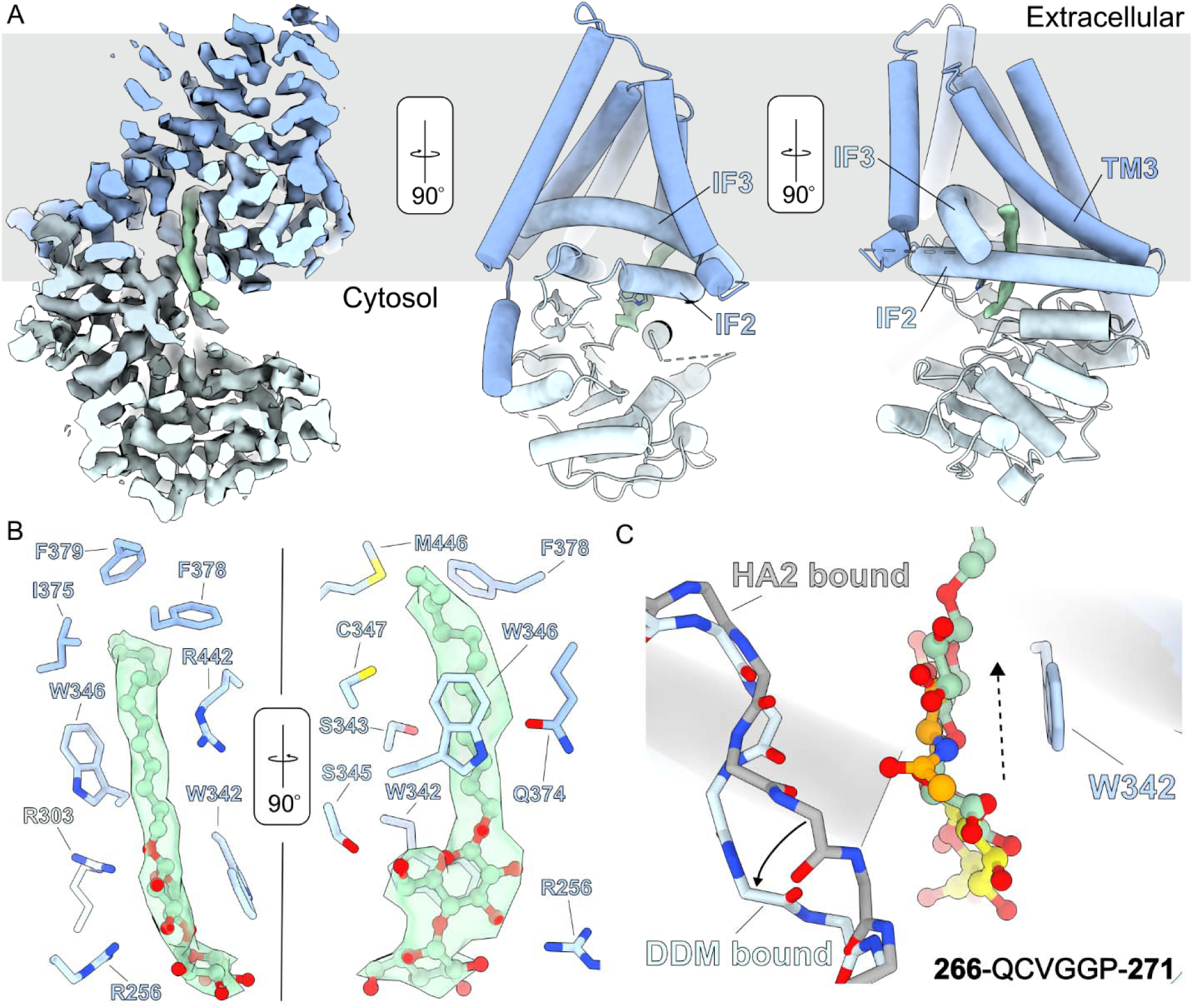
DDM binding at the acceptor site. A, Left - Cross-section of a cryoEM density map for CvHAS (blue) bound to DDM (green). Middle and right – atomic coordinates of CvHAS with DDM cryoEM density independently contoured. Relevant interfacial (IF) and transmembrane (TM) helices are labeled. B, DDM coordination. DDM is shown in ball and stick representation with light green carbon atom coloring. C, Structural alignment of DDM-bound and HA disaccharide (HA2)-bound (PDB ID: 8snc) coordinates. The switch loop (266-271) is displayed with backbone atoms only for clarity, with the DDM-bound conformation shown in light blue and the HA2-bound conformation shown in grey. A solid arrow is used to indicate direction of switch loop flipping from the HA2 bound to DDM bound state. A dashed arrow is used to indicate the direction of displacement of DDM’s maltose group relative to the HA disaccharide.

Relative to the HA disaccharide-bound state, the reducing end glucosyl unit of DDM is shifted toward the TM channel by about 2 Å (Fig 3C). In this position, its C6 hydroxyl group is in hydrogen bonding distance to Ser345, and the ring oxygen interacts with the guanidinium group of Arg303, which is about 3.1 Å away (Fig 3B). Additionally, a previously described ‘switch’ loop (residues 267-271) at the back of the substrate binding groove^8^, moves approximately 3.6 Å away from the maltose moiety relative to the HA disaccharide-bound structure. This conformational change prevents direct interactions of the switch loop with DDM’s head group (Fig 3C and S8).

Strikingly, DDM’s dodecyl alkyl chain extends into a hydrophobic tunnel that is formed by CvHAS’ transmembrane region. The tunnel is formed by Interface Helices two and three as well as TM helix 3 (Fig 3A). It is open to the lipid bilayer environment which likely allows lipid acyl chains to partially enter the enzyme in a biological membrane. The tunnel is lined by Ser343, Trp346, Cys347, Gln374, Ile375, Phe378, Phe379, Arg442 and Met446 (Fig 3B).

### Acceptor promiscuity

The intriguing binding pose of DDM’s maltoside headgroup at the acceptor site prompted us to investigate whether CvHAS accepts other carbohydrates as glycosyl transfer substrates. To this end, we monitored the increase of UDP-GlcA turnover by CvHAS in the presence of selected mono- and disaccharides. As discussed, addition of GlcNAc to a reaction of CvHAS and UDP-GlcA dramatically increases UDP-GlcA turnover, which is monitored using the enzyme-coupled reaction described above.

Under similar conditions, we tested the suitability of the monosaccharides D-galactose, D-glucose, D-mannose, L-rhamnose and L-arabinose to prime CvHAS. None of the monosaccharides tested were able to increase UDP-GlcA turnover above the unsubstituted condition (Fig S9). Next, the disaccharides cellobiose, chitobiose, maltose, sucrose, and xylobiose were tested. Of those, only cellobiose and chitobiose increased UDP-GlcA turnover at elevated concentrations between 10 and 50 mM (Fig 4A). Interestingly, cellotriose and cellotetraose failed to elicit a similar effect on UDP-GlcA turnover (Fig S9).

**Figure 4:**
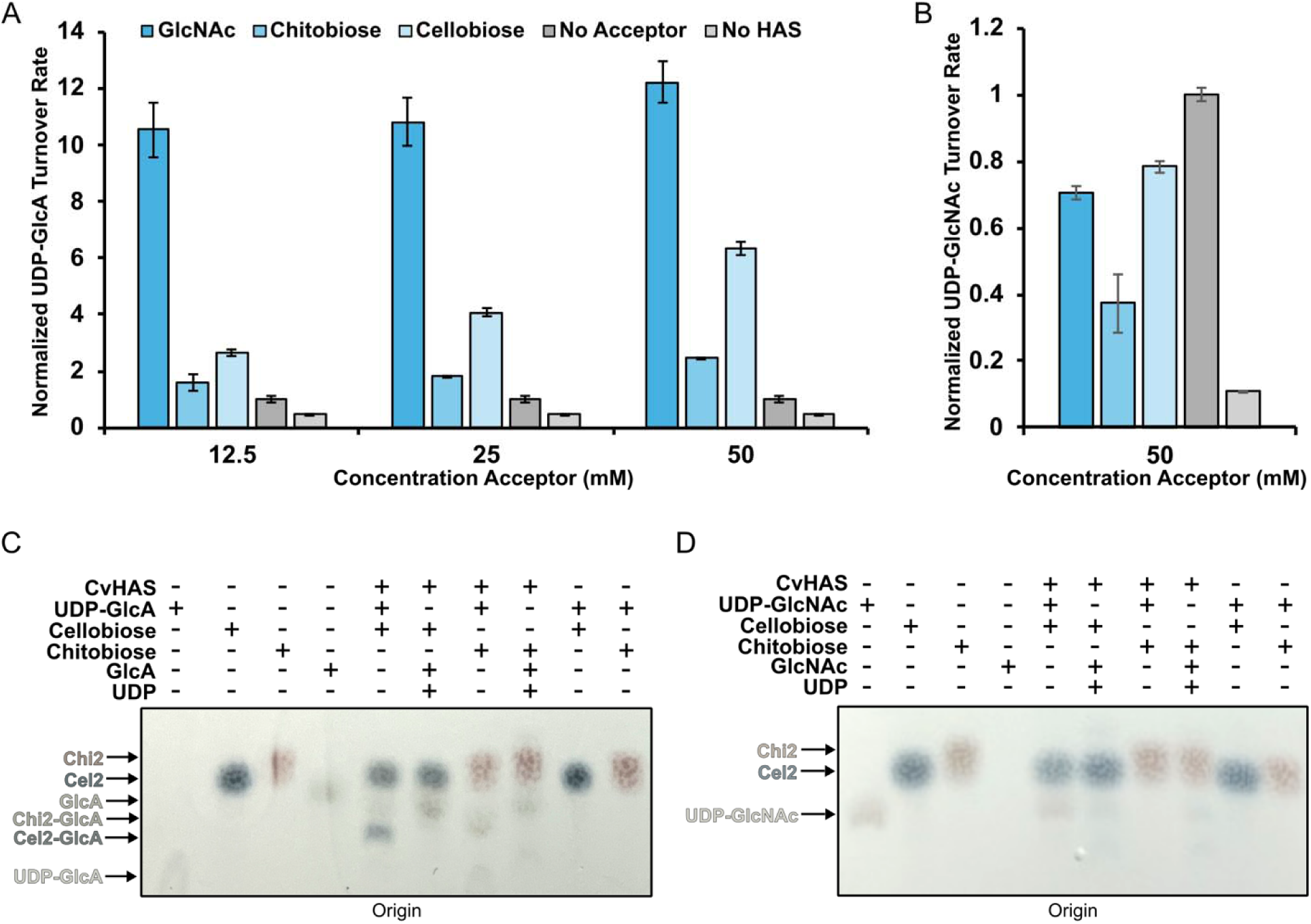
Transfer of GlcA to non-canonical acceptor substrates. A – B, Relative reaction rates for UDP-GlcA (A) and UDP-GlcNAc (B) turnover in the presence of GlcNAc, chitobiose (Chi2) and cellobiose (Cel2). Rates were normalized to samples where the acceptor volume was replaced with water (light grey). Individual velocities were calculated as the average of three technical replicates. Error bars correspond to the standard deviation from the mean. C,D – Diphenylamine staining of TLC plates developed after spotting CvHAS glycosyl transfer reaction mixtures containing either UDP-GlcA (C) or UDP-GlcNAc (D).

In previous studies, GlcA alone failed to prime a UDP-GlcNAc transfer reaction by CvHAS, likely due to instability at the acceptor site ^8^. Alternatively, GlcA stability may be improved when presented in the form of a non-reducing end cap of a HA oligosaccharide. To this end, we tested the effect of titrating a HA tetrasaccharide (HA4) with GlcA at the non-reducing end on UDP-GlcNAc turnover. At up to 25 mM concentration, no apparent change in UDP-GlcNAc turnover rate was observed (Fig. S9), perhaps due to diffusion limitations imposed by a closed HA channel. The results are consistent with previous reports on XlHAS-1 that also failed to elongate an HA tetratsaccharide^18^.

Compared to the GlcNAc monosaccharide, the stimulatory effects of chito- and cellobiose are reduced to about 15 and 50%, respectively, at the highest concentrations tested (50 mM). Consistent with previous observations, no increase in UDP-GlcNAc turnover was observed in the presence of added mono- and disaccharides (Fig 4B). Instead, cello- and chitobiose as well as GlcNAc slightly reduced its turnover, compared to the unsubstituted reaction.

Increasing UDP-GlcA turnover in the presence of cellobiose or chitobiose suggests that the disaccharides may serve as acceptors of the GlcA transfer reaction. We employed thin layer chromatography (TLC) to separate the reactants and products and visualized them with thymol stain (see Methods). As shown in Fig. S10A, UDP-GlcA, GlcA, and cellobiose are appreciably separated on Silica Gel 60 plates using a 5:3:2 volume ration of butanol/ethanol/water as solvent. UDP-GlcA shows the least mobility and remains close to the origin position, clearly separated from GlcA. In the presence of cellobiose and UDP-GlcA, a CvHAS product species can be detected that migrates slightly above the UDP-GlcA position (Fig. S10A).

Repeating the TLC analysis with diphenylamine staining instead of thymol provides a different colorimetric readout for cello- and chitobiose adducts (dark blue and burgundy bands). For both disaccharides, we detect putative monosaccharide adducts under reaction conditions that migrate below the cello- and chitobiose species (Fig 4C). The appearance of these products depends on the presence of CvHAS, UDP-GlcA, and the disaccharide, suggesting that the bands indeed represent novel reaction products of CvHAS. Conversely, replacing UDP-GlcA with UDP-GlcNAc as the substrate did not result in the formation of detectable new products with cello- or chitobiose, suggesting that these acceptors can only be extended with GlcA (Fig 4D).

To confirm that the extended disaccharide species indeed contain GlcA, we repeated the synthesis reaction in the presence of ^14^C-labeled UDP-GlcA. TLC analysis followed by autoradiography revealed species migrating slightly above and below the GlcA monosaccharide band for chito- and cellobiose, respectively (Fig S10C). While the mobility of the individual bands between TLC experiments shows some variability, our data suggests that cellobiose and chitobiose can indeed serve as GlcA acceptors, albeit at high concentrations.

## Discussion

HA biosynthesis is a multi-step process. It involves the sequential binding of UDP-activated GlcA and GlcNAc to HAS’s catalytic site, the transfer of the donor sugar to an acceptor, and the translocation of the nascent HA polymer across the plasma membrane^5^. Important differences exist in how CvHAS interacts with its substrate. The more abundant UDP-GlcNAc substrate is readily bound and hydrolyzed by apo CvHAS to initiate HA biosynthesis. Turnover of the second substrate, UDP-GlcA, however, is inefficient in the absence of an accepting carbohydrate moiety and does not prime the synthesis reaction.

The resolved proofreading and inserted binding poses of UDP-GlcA suggest that recruitment of this substrate occurs in multiple steps. The proofreading pose, in which the carboxylate is recognized between positively charged residues, may serve to distinguish UDP-GlcA from UDP-Glc, which is the more abundant metabolite. Upon its proper insertion into the catalytic cleft, the subtle reorganization of the surrounding priming loop likely positions and stabilizes UDP-GlcA for glycosyl transfer.

Observing two distinct UDP-sugar binding poses is not entirely unprecedented for processive GTs. A comparison of cryo-EM structures for *P. sojae* and *C. albicans* chitin synthase (CHS) in complex with UDP-GlcNAc revealed that CHSs likely utilize an analogous mechanism for substrate insertion^19,20^. In one case, the GlcNAc moiety points almost 180 degrees away from the active site and must therefore undergo large spatial rearrangements to adopt its proper position for glycosyl transfer (Fig S11).

It is currently unclear why GlcA cannot prime HA biosynthesis. The turnover of UDP-GlcA is substantially increased in the presence of an accepting glycosyl unit, likely due to the positioning of a suitable nucleophile to mediate the attack on the donor sugar. Because the apparent dissociation constants for UDP-GlcA in the absence and the presence of GlcNAc are similar, a GlcNAc primer has only modest effects on substrate binding but significantly affects the stability of the substrate at the active site.

Due to exposure during purification, CvHAS accommodates a DDM molecule at its acceptor site and an adjacent hydrophobic tunnel. DDM’s dodecyl tail, which likely mimics a phospholipid acyl chain, seals off a lateral connection to the active site. Coordination of DDM’s reducing end glucosyl unit prompted us to investigate CvHAS’ promiscuity toward alternative primers. Indeed, our findings revealed CvHAS can catalyze unusual biochemistry through the addition of GlcA to cellobiose or chitobiose, albeit at high concentrations. This finding points to degeneracy in the active site of HAS, relaxing specificity toward acceptor substrates. Evolutionary conservation with other QXXRW motif bearing GTs, such as cellulose and chitin synthases, could explain this phenomenon ^21,22^. Both the DDM bound structure, and observations of non-canonical primer extension point to the exploitable potential for HAS and other related GTs to catalyze unnatural carbohydrate synthesis reactions.

Taken together, this work elucidates the molecular basis for dual-substrate selectivity by a model class 1-NR HAS. Visualization of UDP-GlcA in the proofreading conformation establishes a previously uncharacterized checkpoint that supports integrity of HA synthesis. In addition, this study bolsters the previously asserted hypothesis that CvHAS initiates HA synthesis via self-catalyzed formation of a GlcNAc primer through distinguishing acceptor dependent turnover kinetics for UDP-GlcA from UDP-GlcNAc. Findings reported here should inform future investigations of other similar bifunctional GTs, such as the cellulose synthase-like (Csl) enzyme class involved in hemicellulose biosynthesis ^23^.

**Supplementary Figure 1:**
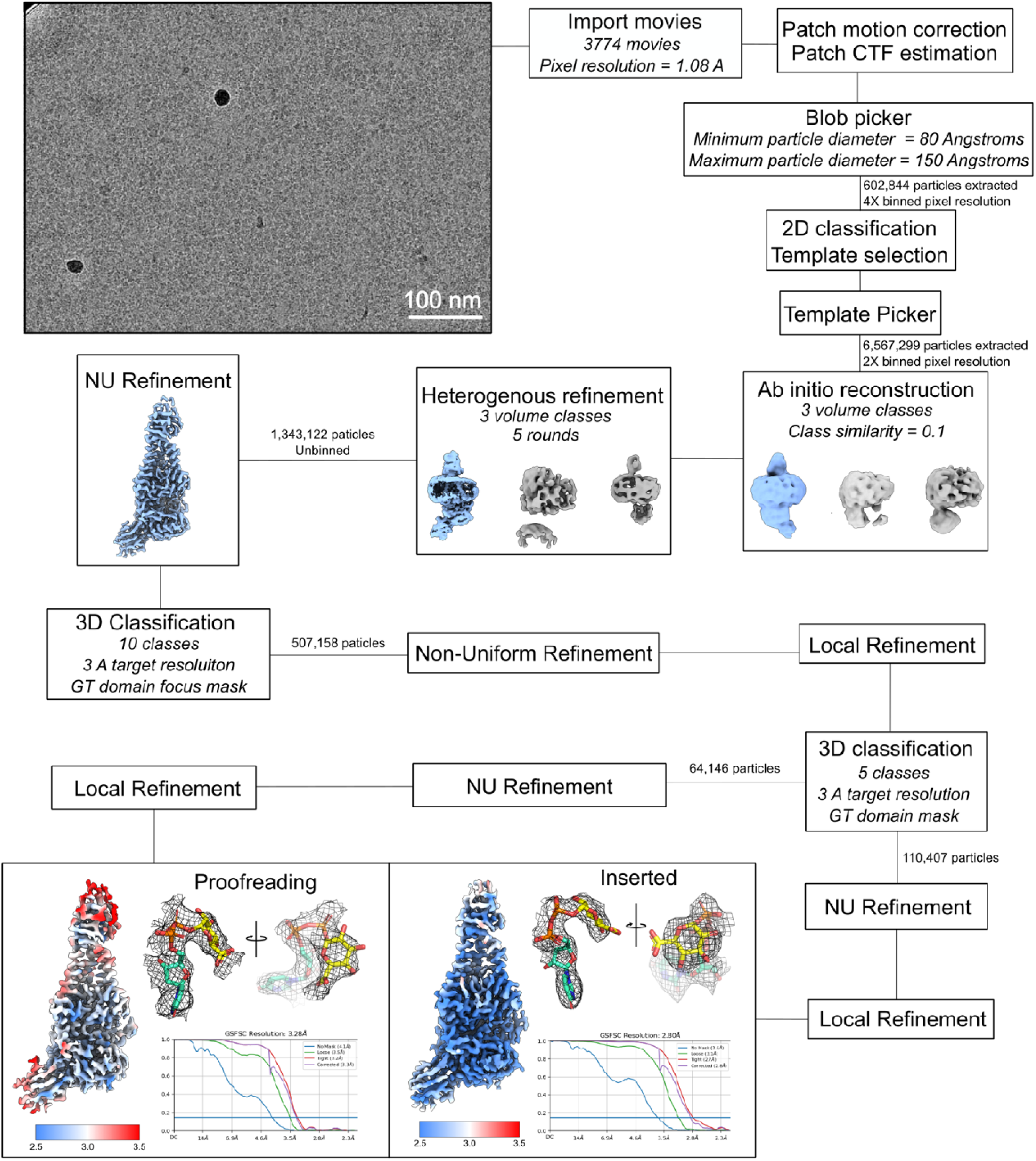
CryoEM data processing for UDP-GlcA inserted and proofreading structures. Carves of cryo electron density maps for UDP-GlcA in inserted and proofreading states are displayed as a black mesh. Local resolution maps are colored according to estimated resolution at FSC = 0.143 in Å.

**Supplementary Figure 2:**
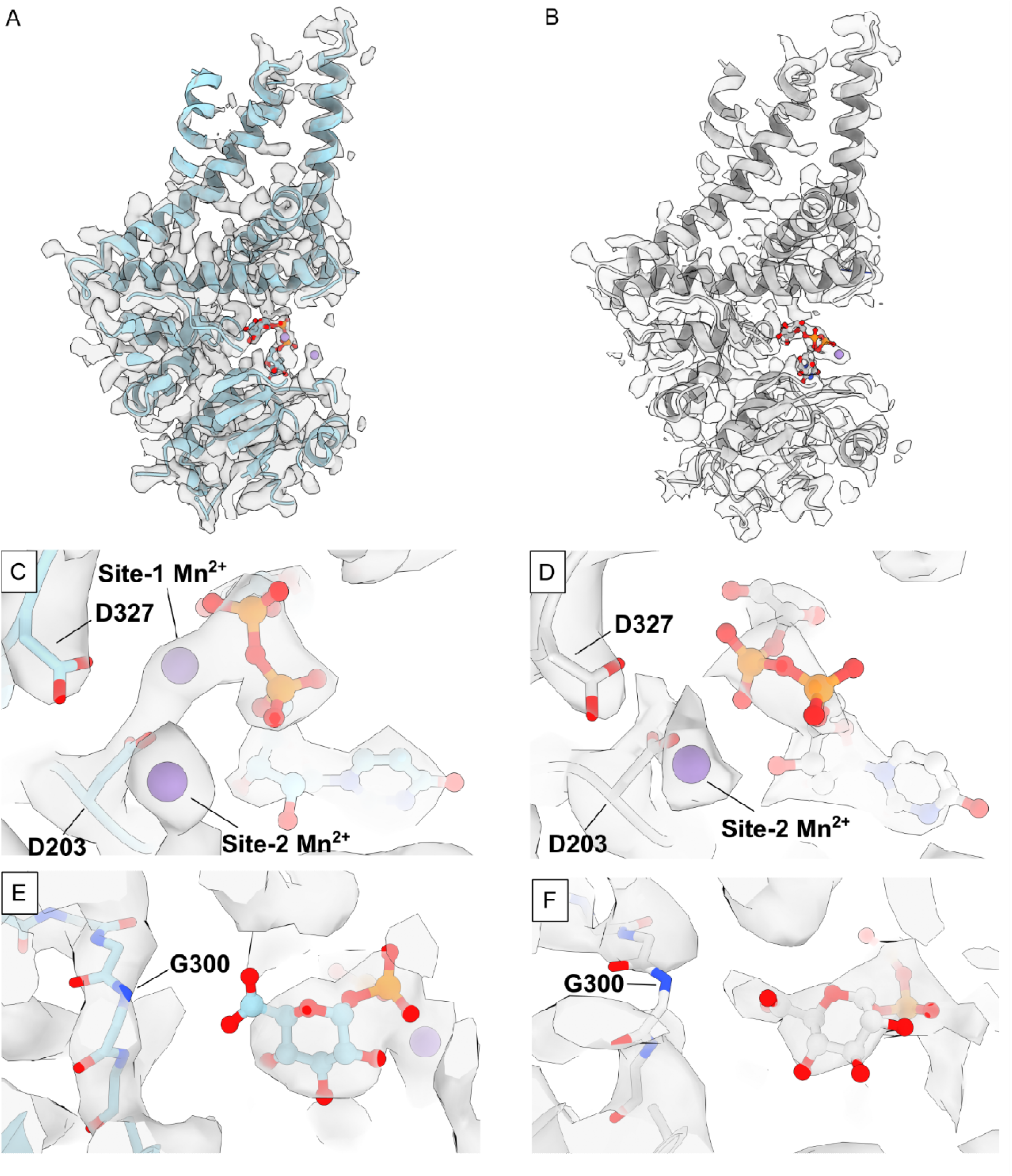
CryoEM density for Mn and the priming loop c-terminus in inserted and proofreading conformations. A, Cross-section of the cryoEM density map for the inserted UDP-GlcA pose. CvHAS and ligand carbon atoms are colored in light cyan, with the ligand represented as a ball and stick model. B, Cross-section of cryoEM density map for the proofreading UDP-GlcA pose. Protein and ligand carbon atoms are colored in grey. C-D Active site view showing local cryoEM density for putative manganese ions (purple) in inserted (C) and proofreading (D) UDP-GlcA bound CvHAS. E, Active site view showing continuous cryoEM density for Gly300 in the inserted UDP-GlcA bound state. F, Broken cryoEM density due to an unresolved Gly300 in the proofreading UDP-GlcA bound state.

**Supplementary Figure 3:**
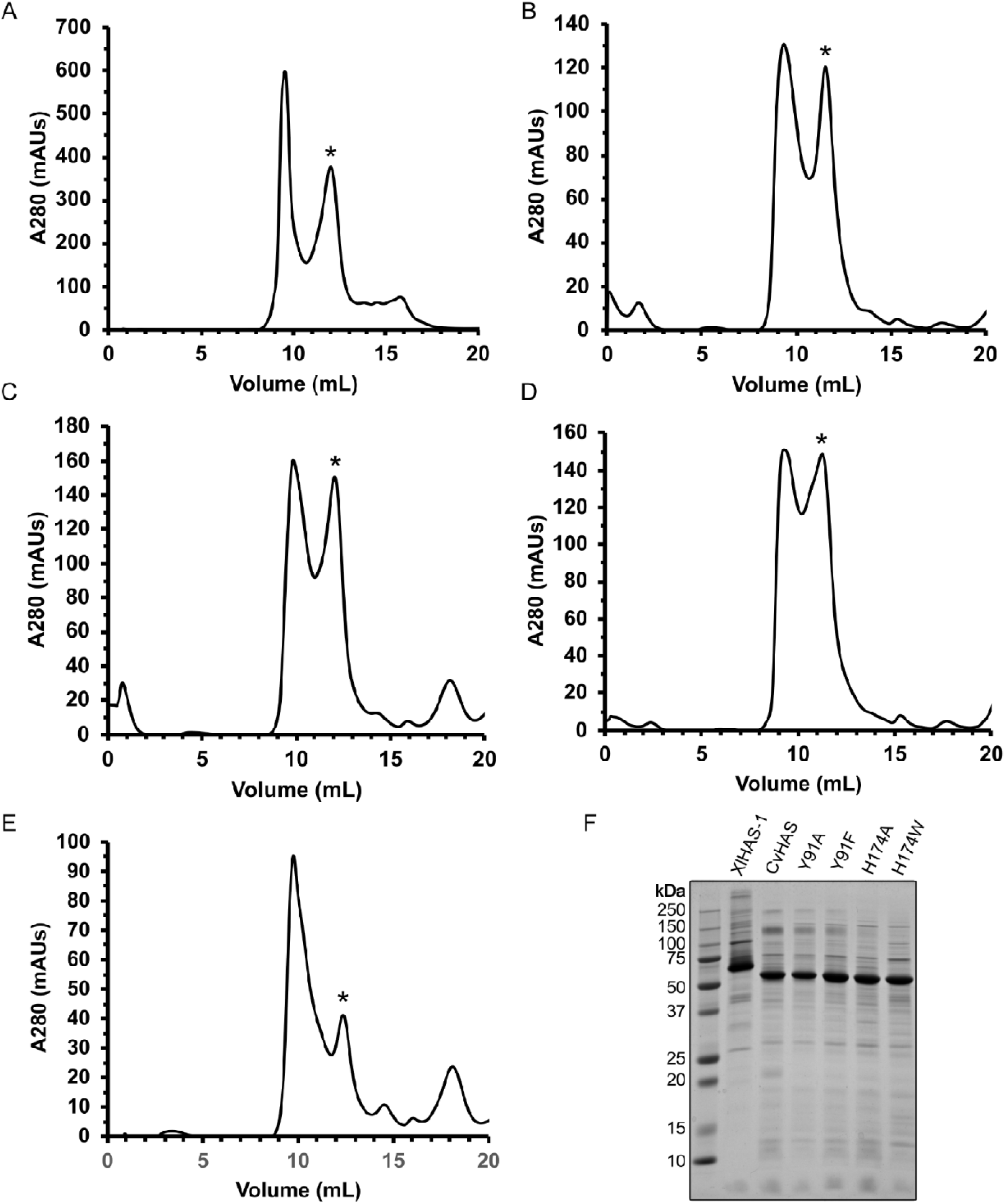
Purification CvHAS’ uracil pocket mutants. A-E, S200 Increase chromatography of WT CvHAS (A), Y91A (B), Y91F (C), H174A (D) and H174W (E). Peaks pooled for subsequent biochemistry are indicated with an asterisk (*). F, Coomassie stained SDS-PAGE gel for purified XlHAS-1, WT CvHAS, and CvHAS mutants.

**Supplementary Figure 4:**
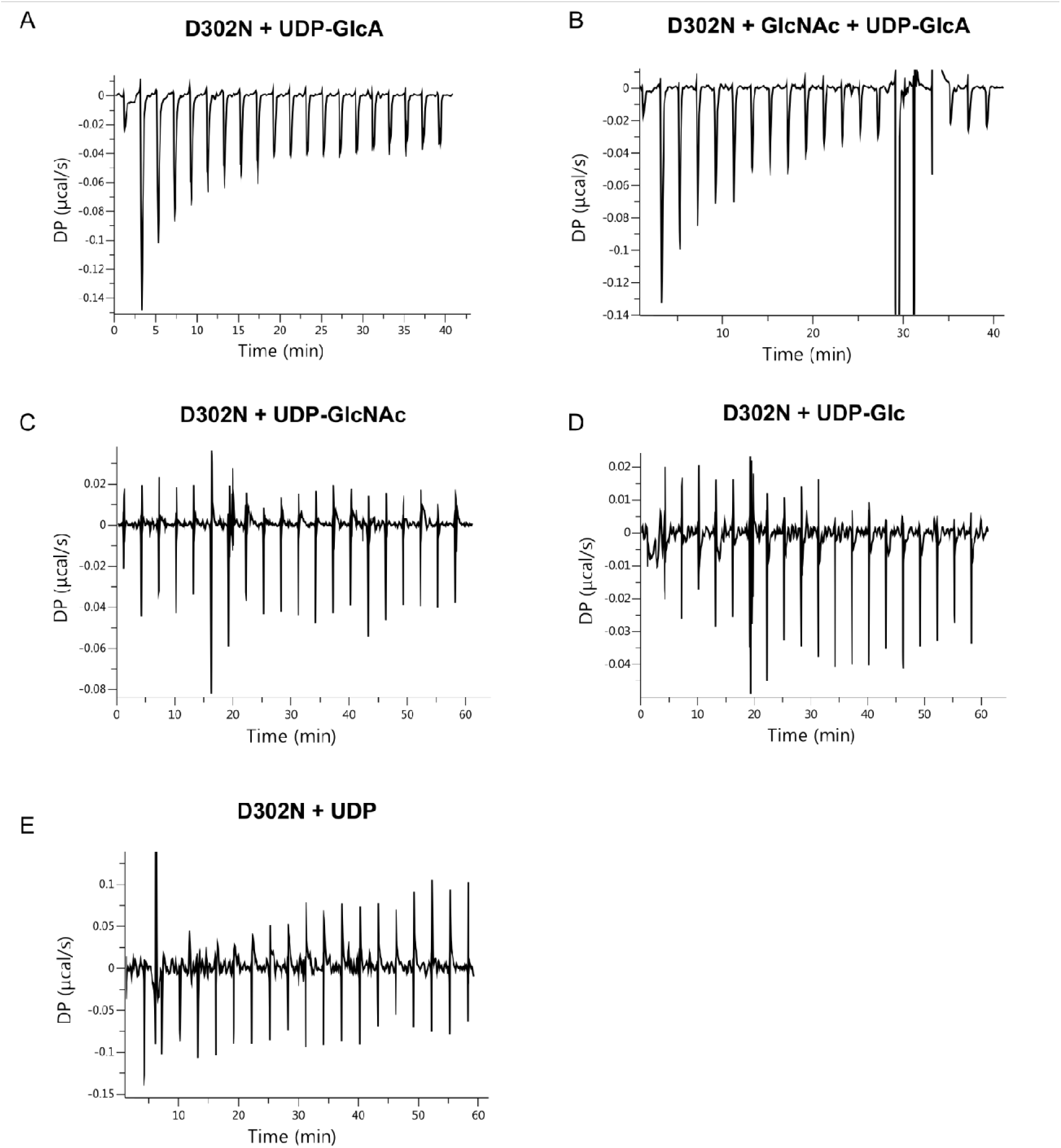
ITC plots for CvHAS substrate titration. A-B, raw ITC plots for UDP-GlcA titration in the absence (A) or presence (B) of excess GlcNAc. C-E, Raw ITC plots for UDP-GlcNAc, UDP-Glc and UDP.

**Supplementary Figure 5:**
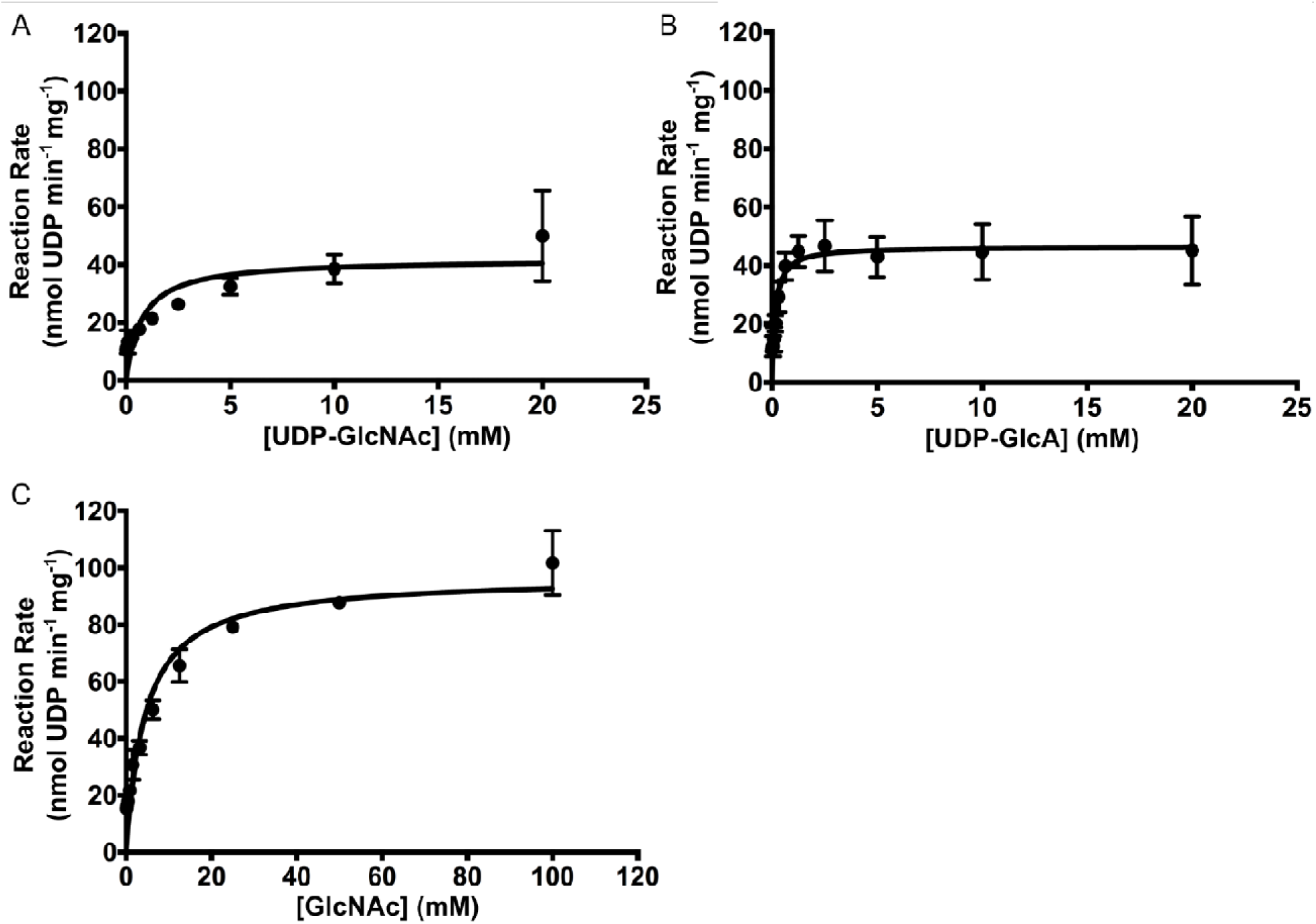
Michaelis-Menten fits for CvHAS substrate titration. A, CvHAS titration with UDP-GlcNAc in the absence of an acceptor. B, Titration of UDP-GlcA in the presence of 10 mM GlcNAc. C, Titration of GlcNAc in the presence of 2.0 mM UDP-GlcA. Five technical replicates were used to derive average reaction rates and standard deviations for plotting of UDP-GlcNAc and UDP-GlcA titrations. Three technical replicates were used for fitting the GlcNAc titration series. All non-linear regressions were performed in Prism 6.0. Error bars represent standard deviations from the means.

**Supplementary Figure 6:**
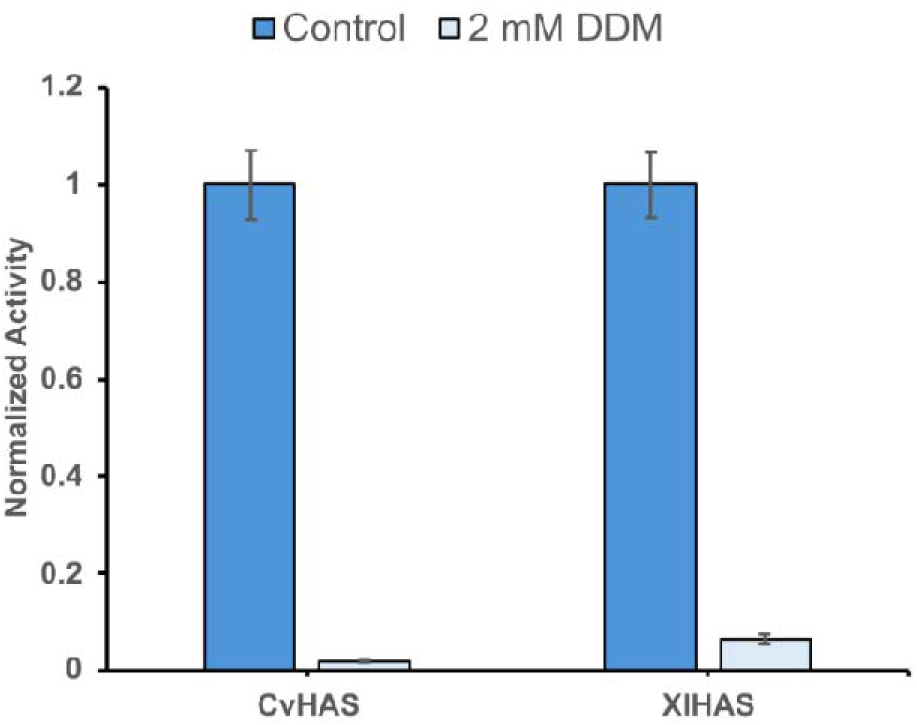
HA synthesis activity is abolished by DDM. Measurements of HAS activity were taken as the average of three technical replicates. Activity values for CvHAS and XlHAS were independently normalized to the control condition. Error bars correspond to the standard deviation from the mean.

**Supplementary Figure 7:**
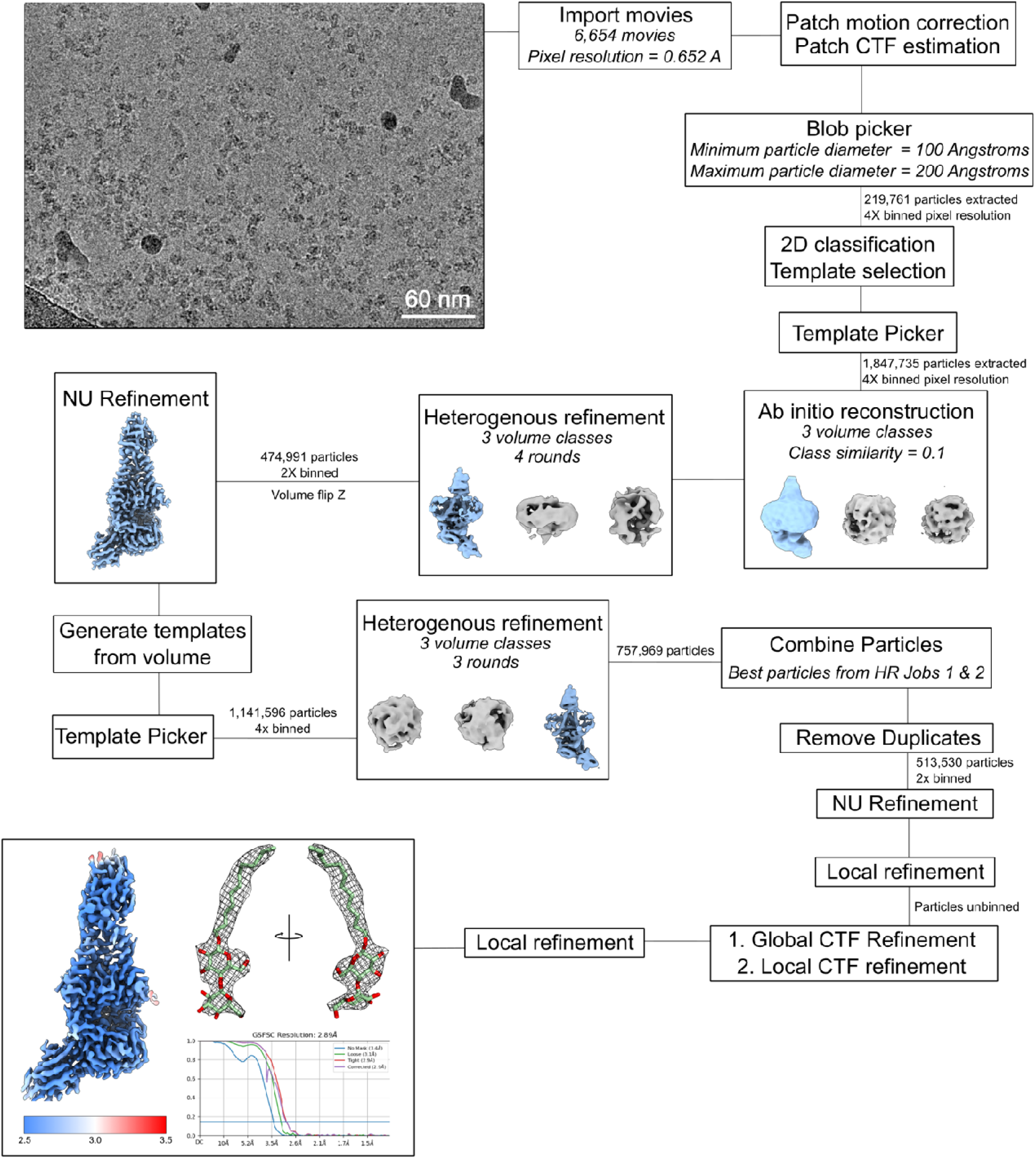
CryoEM Data processing for DDM bound CvHAS. CryoSPARC data processing workflow for CvHAS bound to DDM. Carves of DDM density are shown as a black mesh. Local resolution estimates calculated at FSC = 0.143 are reported in Å.

**Supplementary Figure 8:**
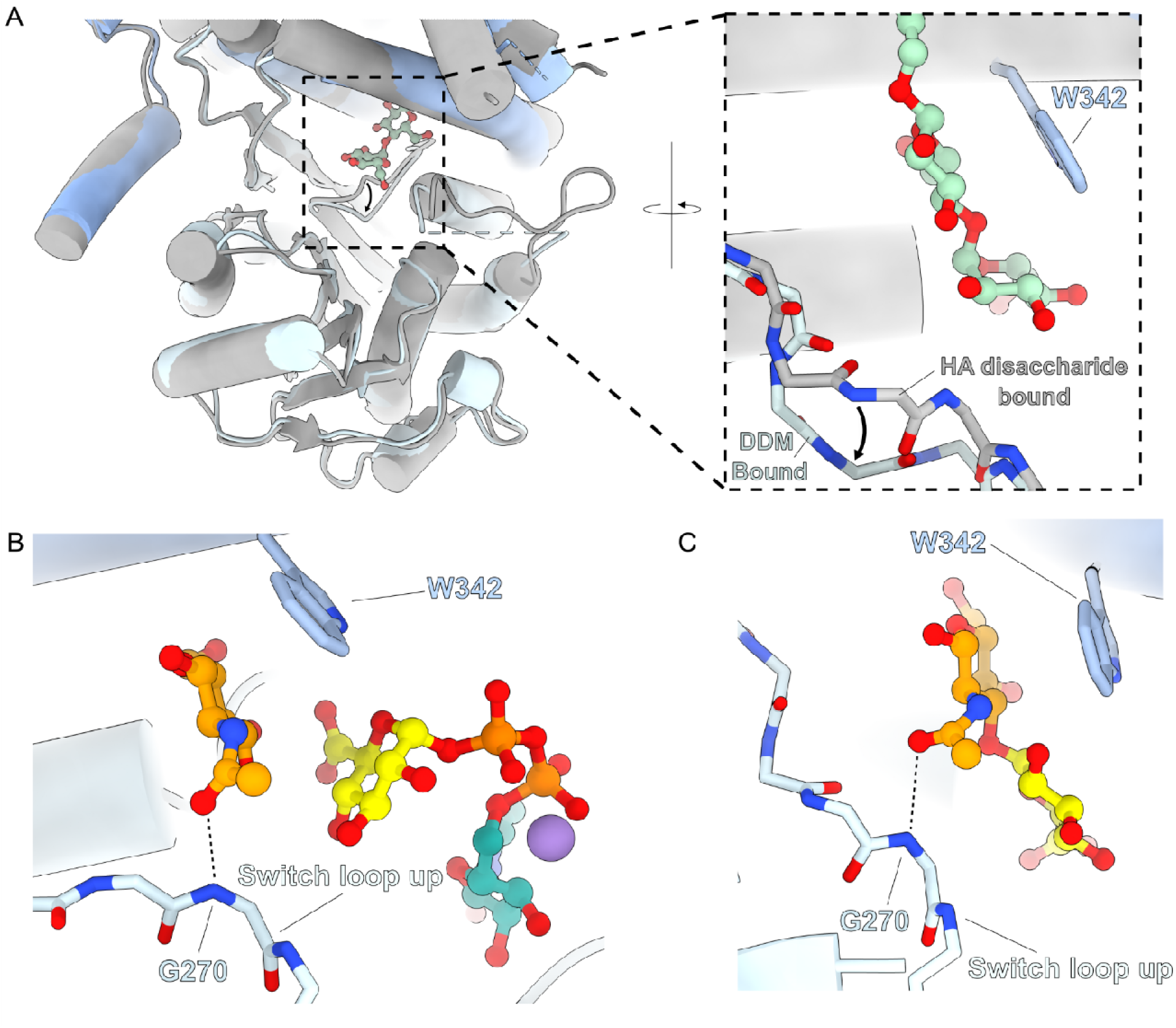
Switch loop movement in GlcNAc, HA disaccharide and DDM-bound CvHAS structures. A, Superimposed structures for the GlcNAc primed, UDP-GlcA-bound CvHAS (grey) and DDM-bound CvHAS (blue). Switch loop movement is indicated by a black arrow. B-C, Position of the switch loop and interactions with GlcNAc in primed, UDP-GlcA-bound (PDB ID: 8snd) CvHAS (B) and HA disaccharide-bound (PDB ID: 8snc) CvHAS (C).

**Supplementary Figure 9:**
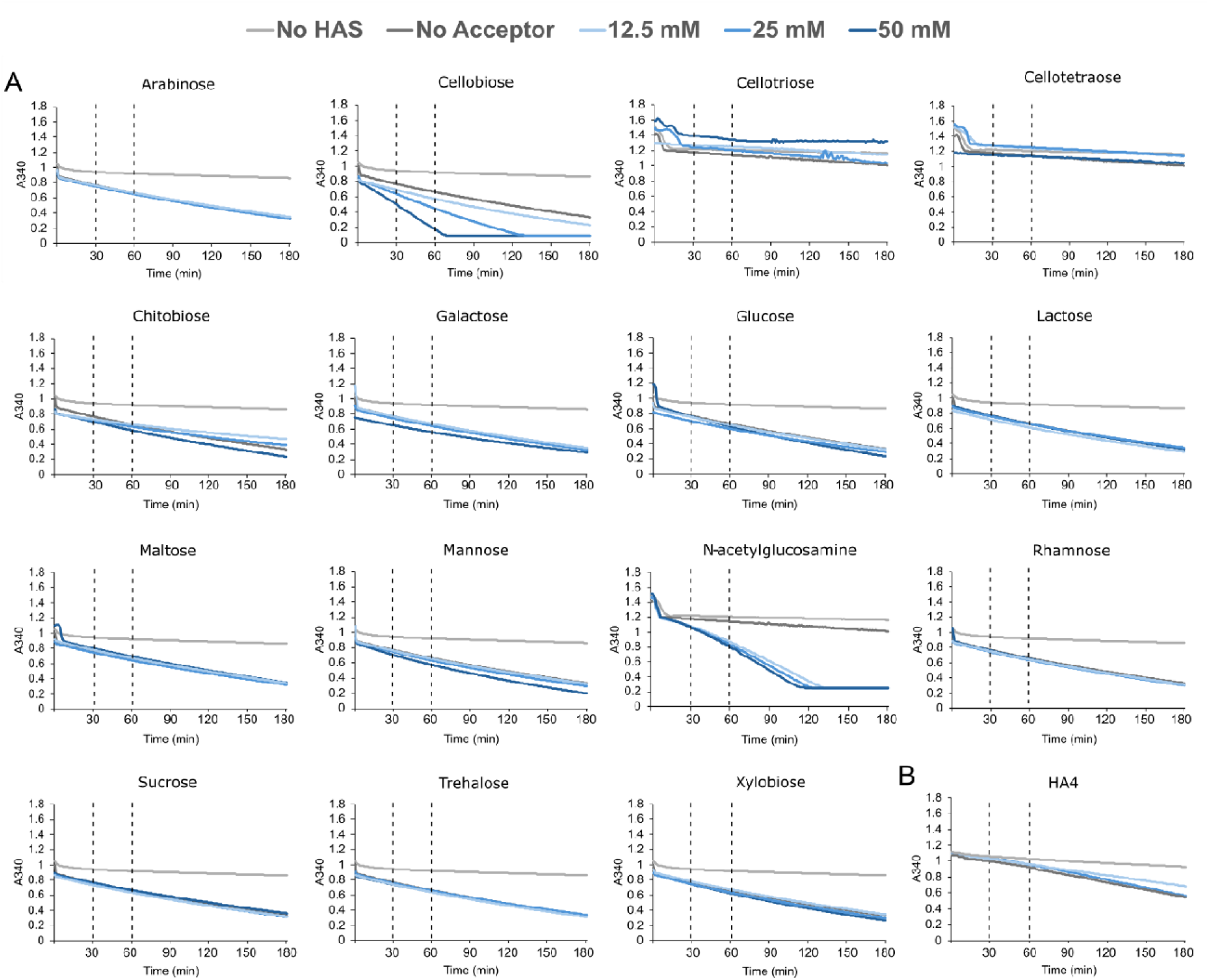
Kinetic traces of UDP-GlcA hydrolysis. Individual kinetic traces for UDP- GlcA (A) and UDP-GlcNAc (B) turnover in the presence of potential glycosyl transfer acceptors supplemented at 12.5, 25, and 50 mM concentrations. The time-interaval used to determine reaction velocities is indicated by two vertical dashed lines.

**Supplementary Figure 10:**
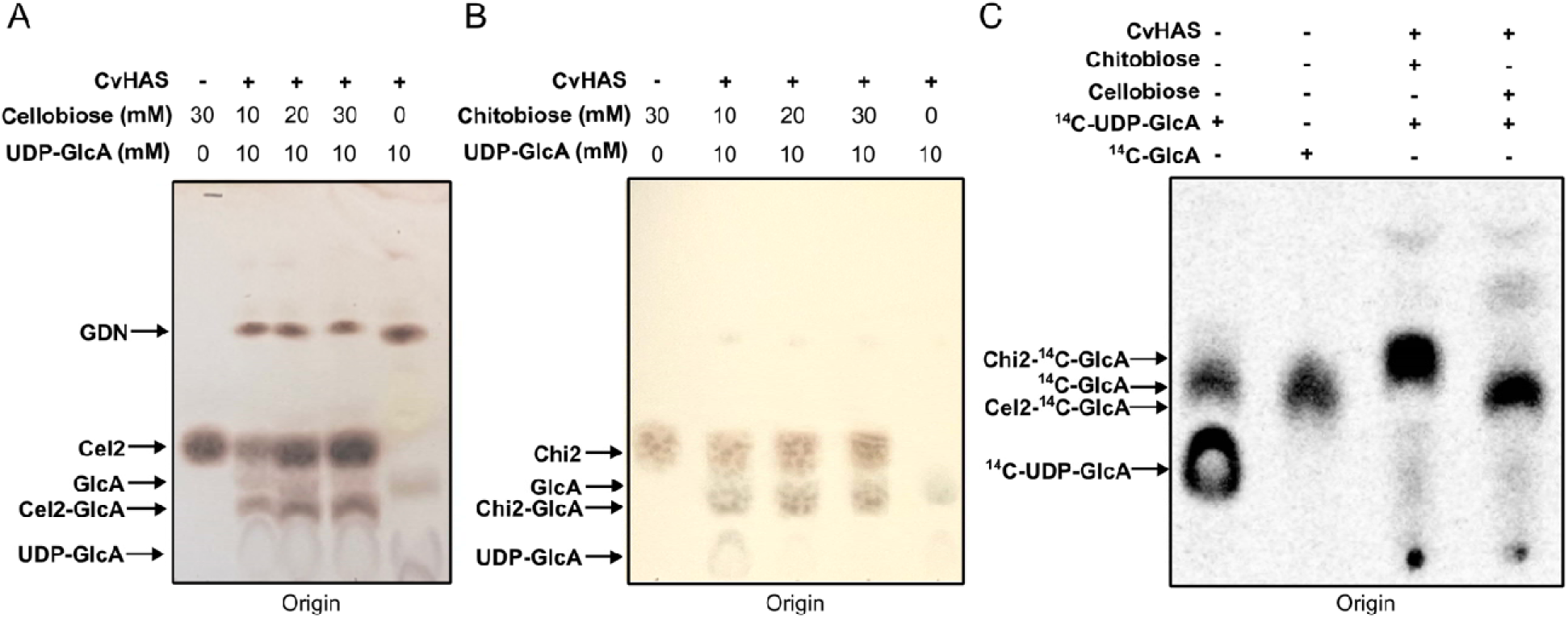
Extension of cellobiose and chitobiose by GlcA. A, TLC analysis of cellobiose (Cel2) titration in the presence of excess UDP-GlcA and CvHAS. Staining was performed with thymol reagent. B, TLC analysis of chitobiose (Chi2) titration. In the presence of excess UDP-GlcA and CvHAS. Staining was performed with diphenylamine reagent. C, Autoradiograph of TLC experiment measuring transfer of ^14^C-GlcA to cellobiose and chitobiose.

**Supplementary Figure 11:**
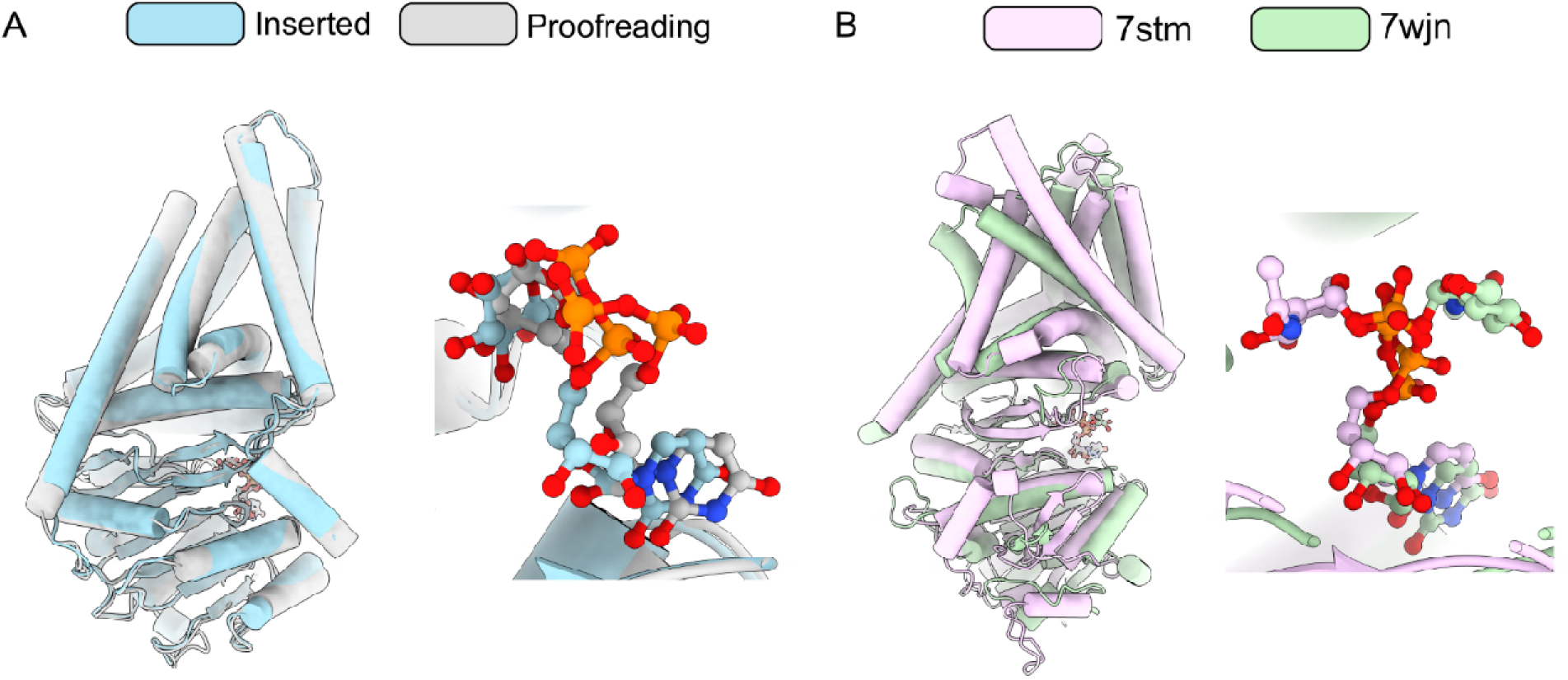
Comparison of UDP-GlcA binding by CvHAS to UDP-GlcNAc binding by CHS. A, Inserted (light cyan) and proofreading (light grey) UDP-GlcA-bound CvHAS structures superimposed. B, Structures of *C. albicans* CHS-2 (PDB ID: 7stm, light purple) and *P. sojae* CHS-1 (PDB ID: 7wjn, light green) bound to UDP-GlcNAc superimposed. 7stm corresponds to the proposed ‘inserted’ UDP-GlcNAc pose for CHS.

**Table 1:** CryoEM data collection, refinement and validation statistics.

|  | **UDP-GlcA, Inserted**<br>**EMDB ID: EMD-73324**<br>**PDB ID: 9YQ5** | **UDP-GlcA, Proofreading**<br>**EMDB ID: EMD-73323**<br>**PDB ID: 9YQ4** | **DDM Bound**<br>**EMDB ID: EMD-73321**<br>**PDB ID: 9YQ2** |
| --- | --- | --- | --- |
| **Data collection and processing** |  |  |  |
| Magnification | 81,000X | 81,000X | 130,000X |
| Voltage (kV) | 300 | 300 | 300 |
| Electron exposure (e- per Å<sup>2</sup>) | 50 | 50 | 60 |
| Defocus range (µm) | -2.0 to -1.0 | -2.0 to -1.0 | -1.8 to -0.8 |
| Pixel size (Å) | 1.08 | 1.08 | 0.652 |
| Symmetry imposed | C1 | C1 | C1 |
| No. of initial particle images | 6,567,299 | 6,567,299 | 6,386,906 |
| No. of final particle images | 110,407 | 64,146 | 513,368 |
| Map resolution (Å) | 2.8 | 3.3 | 2.9 |
| FSC Threshold | 0.143 | 0.143 | 0.143 |
| Map resolution range (Å) | 2.4 - 28.4 | 2.4 - 36.7 | 1.7 - 39.0 |
| **Refinement** |  |  |  |
| Initial model (PDB Code) | 8snd | 8snd | 7sp9 |
| Model resolution (Å) | 2.8 | 3.3 | 2.9 |
| FSC threshold | 0.143 | 0.143 | 0.143 |
| Map sharpening B factor (Å) | 87.5 | 88.8 | 101.8 |
| Model composition |  |  |  |
| Nonhydrogen atoms | 6034 | 5993 | 5939 |
| Protein residues | 742 | 739 | 731 |
| Ligands | 4 | 3 | 4 |
| B-factors (Å<sup>2</sup>) |  |  |  |
| Protein | 93.31 | 45.31 | 55.13 |
| Ligands | 110.36 | 46.85 | 74.2 |
| RMSD |  |  |  |
| Bond Lengths | 0.002 | 0.015 | 0.002 |
| Bond angles | 0.432 | 0.985 | 0.466 |
| Validation |  |  |  |
| MolProbity score | 1.57 | 1.95 | 1.69 |
| Clashscore | 5.36 | 8.34 | 7.29 |
| Poor rotamers (%) | 0.79 | 0.16 | 0.16 |
| Ramachandran plot |  |  |  |
| Favored (%) | 95.91 | 91.66 | 95.84 |
| Allowed (%) | 4.09 | 8.34 | 4.16 |
| Disallowed (%) | 0 | 0 | 0 |

## Methods

### CvHAS Expression and Membrane Harvest

A glycerol stock of *E. coli C43(DE3)* cells harboring a pET28a-CvHAS expression plasmid ^6,8^ was used to inoculate LB broth supplemented with 50 μg mL^-1^ kanamycin and grown overnight. The next day 10 mL of overnight culture was added to 1 L of TB supplemented with 50 μg mL^-1^ kanamycin, 4% glycerol and 1X M salts. Expression cultures were grown to OD600 = 0.8 at 30°C with 220 RPM shaking and cooled to 20°C before induction with 0.5 mM IPTG. Protein expression was allowed to occur overnight before harvesting cell pellets by centrifugation at 4,000 *g* for 10 minutes.

Hereafter, all steps were performed at 4°C unless stated otherwise. Cell pellet taken from 4 L of expression culture was resuspended in RB containing 20 mM Tris-HCl pH 7.5, 100 mM NaCl, 10% glycerol, 0.5 mM TCEP. Lysozyme was added to 1 mg mL^-1^ final concentration and the suspension was mixed for 30 minutes. The cell suspension was disrupted by three rounds of microfluidization at 18,000 PSI, with 1 mM phenylmethysulfonyl Fluoride (PMSF) added after the first passage. Crude lysate was spun at 20,000 *g* for 10 minutes to remove intact cells and larger debris. Supernatant from the first spin was subjected to a second round of centrifugation at 200,000 *g* for 2 hours. The resulting membrane pellet was harvested and flash frozen in liquid nitrogen prior to CvHAS purification.

### CvHAS Purification and Reconstitution

CvHAS purification followed a previously described protocol^8^. Membrane pellet was resuspended in 120 mL of SB containing 20 mM Tris-HCl pH 7.5, 300 mM NaCl, 10% glycerol, 1% DDM, 0.1% cholesteryl hemisuccinate (CHS) and 0.5 mM tris(2-caroboxyethyl)phosphine (TCEP). Subsequently, 1 mM PMSF was added to the membrane suspension and mixed for 1 hour. Following centrifugation at 200,000 *g* for 30 minutes, the supernatant was harvested and batch bound to 5 mL of Ni-NTA resin.

The resin-protein mixture was loaded onto a Kimble flex column and collected by gravity. When purifying CvHAS for functional assays, resin was washed with 20 CVs of WB1 containing 20 mM Tris-HCl pH 7.5, 1M NaCl, 40 mM imidazole, 10% glycerol, 0.05% glycodiosgenin (GDN), 0.5 mM TCEP and 20 CVs of WB2 containing 20 mM Tris-HCl pH 7.5, 300 mM NaCl, 80 mM imidazole, 10% glycerol, 0.02% GDN, 0.5 mM TCEP. CvHAS was eluted in 5 CVs EB (WB2 + 300 mM imidazole). Elutions were concentrated using a 50 kDa MWCO Amicon Ultra Centrifugal Spin Filter (Millipore-Sigma) and injected onto an S200 Increase 10/300 GL (Cytiva) size exclusion column equilibrated in GFB1 containing 20 mM Tris-HCl pH 7.5, 100 mM NaCl, 0.02% GDN, 0.5 mM TCEP.

Nanodisc reconstitution was also performed as previously described^8^. In short, the GDN composition of purification buffers was modified to 0.02% DDM/0.002% CHS. Following size-exclusion, fractions containing CvHAS were concentrated and batch mixed 1:3:3:3:30 with Nb872:Nb881:MSP1E3D1:*E. coli* total lipid extract solubilized in DDM. After 30 minutes, ∼50 mg of hydrated SM2 adsorbant biobeads (Biorad) were added to the mixture. This was repeated once after an additional 30-minute incubation and again after overnight incubation. The reconstitution mixture was cleared of biobeads by filtration, and re-injected on an S200 Increase 10/300 GL column equilibrated in GFB2 containing 20 mM Tris-HCl pH 7.5, 100 mM NaCl and 0.5 mM TCEP. Fractions having CvHAS nanodiscs in complex with Nb872 and Nb881 were identified via Coomassie staining and used in cryoEM sample preparation.

### Nanobody Expression and Purification

*E. coli WK506* harboring a pMES4-Nb expression plasmid^8^ were inoculated from glycerol stocks into LB broth supplemented with 100 μg mL^-1^ ampicillin and 1 mM MgCl_2_ and grown overnight. Two mL of overnight culture was used to inoculate TB supplemented with 100 μg mL^-1^ ampicillin, 1X M salts, 1 mM MgCl_2_, and 0.1% D-glucose. Cells were grown to OD600 = 0.8 at 37°C with 220 RPM shaking, and IPTG was added to a 1 mM final concentration. The shaker temperature was dropped to 27°C, and protein expression was allowed to occur overnight. Cell pellets were harvested the next day by centrifugation at 4,000 *g* for 10 minutes.

Nanobodies were periplasmically extracted by mixing the cell pellet for 30 minutes with hyperosmotic TES buffer containing 20 mM Tris-HCl pH 8.0, 500 mM sucrose, and 0.05 mM EDTA. The extraction mixture was diluted 3-fold in 0.25X TES buffer and mixed for an additional 30 minutes before centrifugation at 200,000 *g* for 30 minutes. The resulting supernatant was batch bound for 1 hour with 5 mL Ni-NTA resin pre-equilibrated in TBS.

Nickel-resin was collected by gravity, washed with 20 CVs Nb-WB1 containing 20 mM Tris-HCl pH 8.0, 1 M NaCl, 20 mM imidazole and with 20 CVs Nb-WB2 containing 20 mM Tris-HCl pH 8.0, 100 mM NaCl, 40 mM imidazole. Nanobodies were eluted in WB2 supplemented with 300 mM imidazole. Nickel-elutions were concentrated over a 10 kDa MWCO centrifugal spin filter prior to injection on an S75 HiLoad size exclusion column equilibrated in Nb-GFB containing 20 mM Tris-HCl pH 7.5, 100 mM NaCl. SEC fractions with nanobodies were pooled and flash frozen for CvHAS reconstitution.

### CryoEM Sample Preparation

For capturing CvHAS in complex with UDP-GlcA, a catalytically inactive mutant of CvHAS with an Asn substitution for Asp302 was used, as described previously^6^. The inactive CvHAS-Nb complex was concentrated to 4.0 mg/mL and supplemented with 10 mM MnCl_2_ and 5 mM UDP-GlcA. The mixture was incubated for 15 minutes on ice, after which 3.0 μL was applied to glow-discharged QF R1.2/1.3 300 mesh Cu grids and blotted for 6 seconds at 4°C/100% humidity. Grids were plunged into liquid ethane using a Mark IV Vitrobot (Thermo Fisher). For the DDM bound map, D302N CvHAS was incubated with 2 mM MnCl_2_ and 1 mM UDP-GlcA for 15 minutes prior to freezing.

### CryoEM Data Collection and Processing

For UDP-GlcA bound structures, all imaging was done on a Titan Krios equipped with a K3 direct electron detector and GIF energy filter at UVA’s Molecular Electron Microscopy Core. Imaging was performed in counting mode at a calibrated pixel resolution of 1.08 A using a 10 eV slit width, with a target defocus range of −2.0 to −1.0 μm and total dose of 50 e^-^/A^2^. Movies were imported to cryoSPARC v4.0.3^24^ for Patch Motion Correction and Patch CTF Estimation. Particles were initially selected by blob picking and extracted as inputs for 2D template generation. Particles from template picking were extracted with a box size of 256 pixels and 2X Fourier cropping. A subset of the initial picks was used to generate three volume references through ab initio reconstruction, converging on one reliable CvHAS-Nb complex volume and two noise volumes. Iterative heterogenous refinement with the ab initio references was used to remove bad picks. The curated particle set was re-extracted at full box size and used for non-uniform refinement of the initial good volume. The refined particles were passed to a 3D classification job with a focus mask covering the GT-domain. Classes containing density for UDP-GlcA were carried over for a second round of 3D classification using the same masking approach. Two classes corresponding to inserted and proofreading states were identified and independently processed using non-uniform and local refinement jobs.

For the DDM-bound structure, imaging of a UDP-GlcA containing sample was performed at a calibrated pixel resolution of 0.652 A. A target defocus range of −0.8 to - 1.8 μm and total dose of 60 e^-^/A^2^ were used. Initial particle picking and curation workflow followed that described for the UDP-GlcA dataset, however particles were extracted at a 400 pixel box size and binned 4X for all steps preceding the first round of non-uniform refinement. Particles were re-extracted at full box size prior to running non-uniform refinement, global/local CTF refinement and local refinement to arrive at a final reconstruction with well-resolved DDM density.

### Model Building and Refinement

An initial model for both inserted UDP-GlcA and UDP-GlcA proofreading structures was taken from PDB ID: 8snd. GlcNAc was deleted manually in COOT^25^. For building coordinates with DDM bound, PDB ID: 7sp9 was selected as the initial model. GlcNAc was deleted from the initial model and DDM (Monomer ID: LMT) was imported and fit into the cryoEM density map in COOT. All three models were iteratively real-space refined in COOT and Phenix^26,27^.

### Isothermal Titration Calorimetry

CvHAS was purified with 10 mM MnCl_2_ supplemented in GFB1. Three hundred μL of 45 μM CvHAS was loaded into the sample cell of a MicroCal PEAQ-ITC (Malvern Panalytical). For injection series without GlcNAc, a syringe concentration of 1 mM UDP-GlcA dissolved in the modified GFB1 was used. The same approach was followed for measuring UDP, UDP-Glc and UDP-GlcNAc binding. For injection series with GlcNAc, 10 mM GlcNAc and 10 mM MnCl_2_ were included in GFB1 for CvHAS purification and UDP-GlcA dissolution. A 20-injection series with 2.0 μL injection volumes and a reference power of 5 μcal/sec was carried out. Data fitting and binding constant (K_d_) derivation was performed using MicroCal PEAQ-ITC analysis software.

### Enzyme Coupled Substrate Turnover Assays

Enzyme-coupled UDP quantification was performed as previously described^6,28^. A stock reaction mix containing 40 mM Tris-HCl pH 7.5, 150 mM NaCl, 20 mM MnCl2, 1 mM TCEP, 1.5 mM NADH, 2 mM phosphoenol pyruvate (PEP) and 2 U/µL Lactate Dehydrogenase/Pyruvate Kinase enzyme cocktail (Sigma) was prepared. A two-fold dilution series beginning at 20 mM for both UDP-GlcNAc and UDP-GlcA was performed across 12 wells of a 96-well black, microclear bottom plate (Grenier). The stock reaction mix was diluted 2-fold with UDP-sugar well solution and 1 μM CvHAS to initiate the reaction. Absorbance was measured at λ = 340 nm each minute over a 1.5-3 hour period at 30°C. Initial reaction velocities were taken as the slope of a linear regression fit to data points between 30-60 minutes of each kinetic trace. Non-linear Michaelis-Menten fitting was performed with Prism 6.0. When measuring the influence of alternative glycosyl transfer acceptors on substrate turnover, UDP-GlcA or UDP-GlcNAc was included in the stock reaction mix at 4.0 mM. Concentrations of monosaccharides and disaccharides used for screening were varied between 12.5-50 mM.

### HA Gel Electrophoresis

A 1% ultrapure agarose gel (Sigma) was cast with 1X TAE buffer. Reactions were setup as described for substrate hydrolysis measurements and quenched after 1.5 hours with Laemmeli buffer. Agarose gel electrophoresis was run at 100V for 2 hours in 1X TAE. The resulting gel was fixed in 50% EtOH for 1 hour, and subsequently placed in 0.005% Stains-All dissolved in 50% EtOH overnight under dark. The stained gel was transferred to 20% EtOH solution and again left overnight. The next day the gel was exposed to ambient light to remove remaining Stains-All background prior to imaging, as described^11^.

### HA Quantitation by Paper Chromatography and Liquid Scintillation Counting

Radiometric HA quantification was performed as described^17^. HA synthesis reactions were initiated by mixing 10 μM CvHAS 1:1 with a reaction mix containing 80 mM Tris-HCl pH 7.5, 150 mM NaCl, 1 mM TCEP, 40 mM MnCl_2_, 10 mM UDP-GlcA, 10 mM UDP-GlcNAc, 0.1 μCi ^3^H-UDP-GlcNAc (Revvity). Reactions were allowed to occur for 2 hours at 30°C before quenching with 2% SDS. Samples were spotted on Whatman filter paper and dried. The paper was developed in a 65% 1 M ammonium acetate / 35% EtOH mobile phase for 2 hours and dried. The origin was extracted for liquid scintillation counting in a Hidex 300 SL by exposure to UltimaGold scintillation fluid (Revvity).

### TLC of CvHAS Reaction Products

Reactions for thin layer chromatography (TLC) were carried out by mixing 10 μM CvHAS 1:1 with 80 mM Tris-HCl pH 7.5, 150 mM NaCl, 20 mM MnCl_₂_, 1.0 mM TCEP and 10 mM UDP-GlcA, followed by incubation at 30°C for 2 hours. To observe acceptor extension, cellobiose and chitobiose were included in the reaction mix at 10-30 mM final concentration. Each reaction was mixed 1:1 with 50% MeOH, and 2.0 μL was spotted on a Silica Gel 60 plate. TLCs were developed in a *n*-butanol/ethanol/water (5:3:2, v/v/v) solvent system, dried and stained by exposure to either 0.5% thymol (w/v) dissolved in 50:1 EtOH/H_2_SO ^29,30^ or 16.7% diphenylamine (w/v) dissolved in 4:3:17 aniline/H_3_PO_4_/acetone^31^. For analysis of radioactive products, 0.02 μCi of ^14^C-UDP-GlcA was included in the reaction mix. Autoradiography was performed by exposing the TLC plate to a phosphor screen for two days prior to phosphor imaging on a Typhoon IP instrument (Amersham).

### Site Directed Mutagenesis

Complementary forward and reverse primers with the integrated mutant codon annealing at the mutation site, as well as forward and reverse primers for the T7 promoter and T7 terminator sequence of the pET28a-CvHAS vector were generated (IDT). For each mutation, three PCRs amplifying the region of the plasmid sequence from T7 promoter to mutation site, mutation site to T7 terminator and T7 terminator to T7 promoter were carried out in parallel. Resulting amplicons were purified by gel-extraction (Qiagen) and used in a HIFI reaction (NEB) for plasmid assembly. The assembly reactions were transformed into chemically competent DH5α cells, followed by DNA purification and full-plasmid sequence verification.

## Acknowledgements

We are grateful to Michael Purdy and David Cooper at the Molecular Electron Microscopy Core facility for help and support in cryo-EM data collection. The project was in part funded by NIH grant R35GM144130. J.Z. is an Investigator of the Howard Hughes Medical Institute. This article is subject to HHMI’s Immediate Access to Research policy, which requires that this article be made publicly available as initial and revised preprints deposited on a designated preprint server under a CC BY 4.0 license.

## Author contributions

J.Z., Z.S., and J.K. designed the experiments. J.K. performed TLC experiments and Z.S. performed all other biochemical and structural biology procedures. All authors evaluated and interpreted the data. Z.S. and J.Z. wrote the manuscript and all authors edited it.

## Data Availability Statement

Cryo-EM maps have been deposited in the EMDB under the accession codes EMD-73321, EMD-73323 and EMD-73324. Protein coordinates have been deposited in the PDB under the accession codes 9YQ2, 9YQ4 and 9YQ5.

